# Pore-C Simultaneously Captures Genome-wide Multi-way Chromatin Interaction and Associated DNA Methylation Status in Arabidopsis

**DOI:** 10.1101/2022.01.20.477161

**Authors:** Zhuowen Li, Yanping Long, Yiming Yu, Fei Zhang, Hong Zhang, Zhijian Liu, Jinbu Jia, Weipeng Mo, Simon Zhongyuan Tian, Meizhen Zheng, Jixian Zhai

## Abstract

In the past decade, genome-wide characterization of the three-dimensional chromatin structure in plants using high-throughput methods has greatly advanced our knowledge in plant genome architecture (Liu and Weigel, 2015; Ouyang et al., 2020). However, due to the limitation of Illumina short-read sequencing, the genome-wide contact map obtained by Hi-C/ChIA-PET is pairwise, and the multi-way interaction can only be inferred from the two-way data. To directly capture multi-way interaction in Arabidopsis, we applied a long-read-based method called Pore-C that directly sequences the DNA fragments joined by proximity-based ligation (Ulahannan et al., 2019).

---

Compared with Hi-C, Pore-C replaced the Illumina short-read sequencing with Nanopore long-read sequencing, which drastically simplified the library construction procedure by removing the biotin labeling, fragmentation and several affinity-purification and DNA purification steps (Figure 1A). *In-situ* ligation makes Pore-C reads to retain the interaction information within the individual nucleus. We generated two Pore-C libraries using 12-day-old Arabidopsis seedlings of wildtype Col-0 and analyzed the data according to the Pore-C pipeline (Ulahannan et al., 2019). We obtained 6.5 million (11.7 Gb) and 3.5 million (7.9 Gb) high-quality reads (Q7 filter) with N50 up to 2654 bp and 3006 bp, respectively (Supplemental table 1). More than 96% of the reads can be mapped to the Arabidopsis TAIR10 reference genome in both libraries. From about 10 million Pore-C reads in total, we obtained 50.7 million valid pairs of interaction, a similar amount but with fewer reads compared to previous Hi-C data (59.1 million pairwise contacts from 239.5 million Illumina reads) (Wang et al., 2015), demonstrating the efficiency of long-read sequencing in harvesting chromosomal interaction. From the 2D-contact matrixes, the two Pore-C libraries and the Hi-C data showed high consistency in their contact feature, suggesting high reproducibility and accuracy of Pore-C (Figure 1B, C).

**Figure1.**
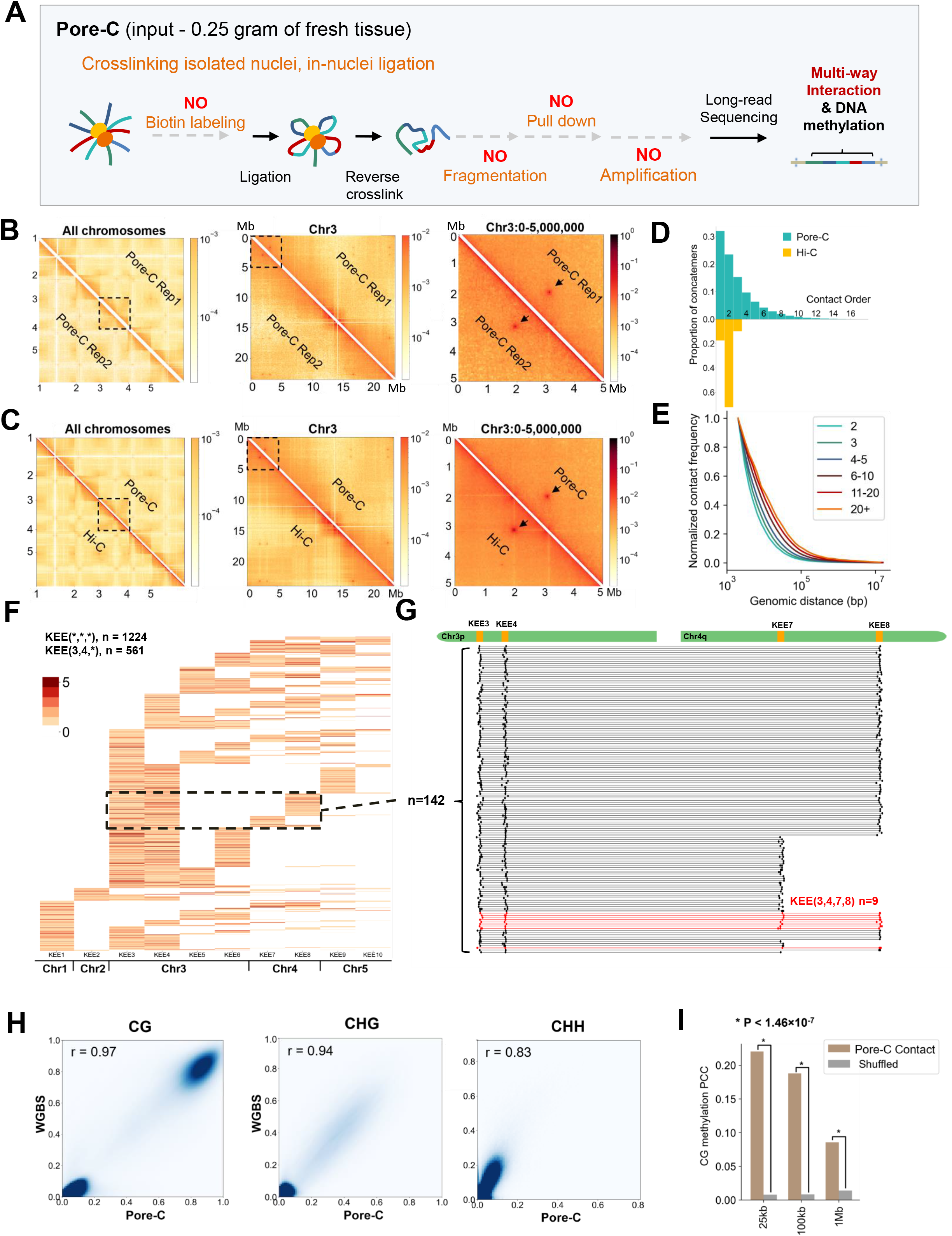
Pore-C captures multi-way contacts and DNA methylation in Arabidopsis. (A) Schematic of the Pore-C protocol. (B,C) The consistency between two Pore-C libraries (B), between Hi-C and Pore-C (C), respectively. Left: contact maps across all chromosomes. Middle: contact maps on chromosome 3. Right: contact maps within chr3: 0-5 Mb. The interactions between KEEs are marked by black arrows. (D) Comparison of contact order distribution between previous Hi-C data and Pore-C data in Arabidopsis. (E) Contact frequency between different genomic distances of various order groups of Pore-C. (F) Combination information of reads related to at least three different KEEs. Each row of the heatmap represents one read. The color intensities in the heatmap indicate the counts of interaction capture within one KEE on one read. KEE(*,*,*) means the interaction between at least three individual KEEs, “*” indicates at least one KEE, and “n” shows the read counts. (G) Alignment locations of individual reads related to KEE3, KEE 4, KEE7 and KEE8. The red reads (n=9) show the direct interactions between all the four KEEs. (H) The consistency of CG, CHG, CHH DNA methylation between Pore-C library and WGBS data. (I) Pearson correlation coefficient (PCC) of CG methylation level between Pore-C contacts with genomic distance larger than 25 kb,100 kb and 1 Mb, respectively.

The Pore-C pipeline defined the number of genome-matched fragments on a Pore-C read as the “Contact Order” (Ulahannan et al., 2019). Our results show that 44% reads contain multi-way interactions (Contact Order ≥ 3), similar to Pore-C in human cells (47.56%) (Ulahannan et al., 2019). Whereas in the Hi-C data (Wang et al., 2015), the Contact Orders are mostly at two, consistent with its design (Figure 1D). Pore-C results in Arabidopsis and animals show a higher efficiency in capturing multi-way interaction compared to SPRITE (12%) (Quinodoz et al., 2018), demonstrating an advantage of the long-read approach over the labeling approach in capturing multi-way interaction. The contact frequency in relation to the genomic distance for our Pore-C data showed that reads with higher contact order detected interactions across longer genomic regions (Figure 1E), which can also give help in *de novo* genome assembly.

To validate the effectiveness of Pore-C in multi-way interaction in Arabidopsis, we examined the interactions among KNOT ENGAGED ELEMENT (KEE) (Grob et al., 2014) with our Pore-C data. In Arabidopsis, the interaction of ten KEEs (named KEE1 to KEE10) can be visualized as discrete dots on the pairwise contact matrix of Hi-C and their pairwise interactions were confirmed by light microscopy-based FISH assay (Grob et al., 2014). However, it remains unclear if multiple KEEs can simultaneously interact together in the same nucleus. In our Pore-C data, 1145 reads simultaneously detect contact among three KEEs, higher than that of the 100 randomly selected control regions (mean 279.29 reads, *P*-value = 0.04), and 75 reads capture the contact among four KEEs and more (Figure 1F). Interestingly, nearly half of these high-order interactions involve both KEE3 and KEE4 (561 reads), suggesting that these two KEEs serve as key hubs in the KEE interacting network (Figure 1F). Direct interaction among four KEEs was also detected, including 9 reads for KEE3, KEE4, KEE7 and KEE8 (referred to as KEE(3,4,7,8), Figure 1G), 4 reads for KEE(1,3,4,6), and 11 reads for KEE(3,4,5,6). These results demonstrate the existence of multi-way contacts of KEEs.

The PCR-free strategy of Pore-C enables to directly detect DNA methylation states and higher-order chromatin interaction by long-read sequencing and thus helps to reveal their coordination on the same read, without the requirement of bisulfite conversion in the Methyl-HiC method (Li et al., 2019). The CG, CHG and CHH methylation level we called from Pore-C reads are highly consistent with WGBS (whole-genome bisulfite sequencing), the gold standard for DNA methylation (Figure 1H) (Ni et al., 2021; Stroud et al., 2012). We used nanopolish (Simpson et al., 2017) to detect 5mCG related to each Pore-C read, and found a higher correlation of CG methylation level among interacting fragments in Arabidopsis (Figure 1I), which is consistent with the previous report in mouse ESCs (Li et al., 2019).

In summary, Pore-C efficiently captures the genome-wide multi-way chromatin interaction landscape at a single-molecular level, and can reveal the epigenetic modification and chromatin organization of the same group of interacting DNA fragments, which may help to explore the heterogeneity of the chromatin interaction and DNA methylation in nuclei of different cell types. With this method, we validated the multi-way contacts among KEEs within nucleus, and found the CG methylated regions on the genome tends to contact. Taken together, our results demonstrate that Pore-C is a simple, accurate and effective method for exploring multi-way chromatin interaction in plants.

## Acknowledgments

Group of J.Z. is supported by the National Key R&D Program of China Grant(2019YFA0903903), the Program for Guangdong Introducing Innovative and Entrepreneurial Teams (2016ZT06S172), the Shenzhen Sci-Tech Fund (KYTDPT20181011104005) and a Stable Support Plan Program of Shenzhen Natural Science Fund Grant (20200925153345004), and Key Laboratory of Molecular Design for Plant Cell Factory of Guangdong Higher Education Institutes (2019KSYS006).

## Author Contributions

Y.L. and Z.L. developed the method and performed the experiments. Z.L., Y.Y., F.Z. and Y.L. analyzed the data. H.Z. set up the website, J.Z. conceived and oversaw the study. Z.L., Y.L. and J.Z. wrote the manuscript. All authors revised the manuscript.

## Materials and Methods

Please refer to Supplementary material.

## Data availability

The Pore-C data generated can be found in the Genome Sequence Archive in National Genomics Data Center under accession number CRA005105 (reviewer link: https://ngdc.cncb.ac.cn/gsa/s/INkjuqHu.

## References

Grob, S., Schmid, M.W. and Grossniklaus, U. (2014) Hi-C analysis in Arabidopsis identifies the KNOT, a structure with similarities to the flamenco locus of Drosophila. Molecular cell 55, 678–693.

Li, G., Liu, Y., Zhang, Y., Kubo, N., Yu, M., Fang, R., et al. (2019) Joint profiling of DNA methylation and chromatin architecture in single cells. Nature methods 16, 129 991–993.

Liu, C. and Weigel, D. (2015) Chromatin in 3D: progress and prospects for plants. Genome biology 16, 170.

Ni, P., Huang, N., Nie, F., Zhang, J., Zhang, Z., Wu, B., et al. (2021) Genome-wide detection of cytosine methylations in plant from Nanopore sequencing data using 134 deep learning. bioRxiv, 2021.2002.2007.430077.

Ouyang, W., Xiong, D., Li, G. and Li, X. (2020) Unraveling the 3D genome architecture in plants: present and future. Molecular Plant 13, 1676–1693.

Quinodoz, S.A., Ollikainen, N., Tabak, B., Palla, A., Schmidt, J.M., Detmar, E., et al. (2018) Higher-order inter-chromosomal hubs shape 3D genome organization in 139 the nucleus. Cell 174, 744–757.e724.

Simpson, J.T., Workman, R.E., Zuzarte, P.C., David, M., Dursi, L.J. and Timp, W. (2017) Detecting DNA cytosine methylation using nanopore sequencing. Nature 142 methods 14, 407–410.

Stroud, H., Hale, C.J., Feng, S., Caro, E., Jacob, Y., Michaels, S.D., et al. (2012) DNA methyltransferases are required to induce heterochromatic re-replication in Arabidopsis. PLoS genetics 8, e1002808.

Ulahannan, N., Pendleton, M., Deshpande, A., Schwenk, S., Behr, J.M., Dai, X., et al. (2019) Nanopore sequencing of DNA concatemers reveals higher-order features of chromatin structure. bioRxiv, 833590.

Wang, C., Liu, C., Roqueiro, D., Grimm, D., Schwab, R., Becker, C., et al. (2015) Genome-wide analysis of local chromatin packing in Arabidopsis thaliana. Genome research 25, 246–256. 152

