## Supplemental table 1 for "Pore-C Simultaneously Captures Genome-wide Multi-way Chromatin Interaction and Associated DNA Methylation Status in Arabidopsis"

**Materials and Methods**

**Plant materials**

The wildtype Arabidopsis seedlings (Col-0) used for constructing Pore-C library were grown on 1/2 MS plates at 22 °C (16 h light/8 h dark) and collected after 12 days.

**Pore-C library construction**

The Pore-C libraries were generated by modifying the previously described Arabidopsis Hi-C and human Pore-C protocol (Feng et al., 2014; Grob et al., 2014; Ulahannan et al., 2019). In brief, ~0.25 gram tissue was ground to fine powder in liquid nitrogen and transferred to 1 ml Extraction Buffer I (0.4 M sucrose, 10 mM Tris-HCl (pH 8.0), 10 mM MgCl_2_, 0.2% Triton X-100, 1 mM EDTA (pH 8.0), 5 mM β-Mercaptoethanol, 0.1 mM PMSF). The homogenate was crosslinked with 133 μl 16% formaldehyde (final is 1%) for 15 min with rotation at room temperature. Then 125 μl 2 M glycine was added and incubate for another 5 min to stop crosslink (final is 0.125 M). After filtering through two layers of miracloth, the lysate was loaded on the surface of 2 ml dense sucrose buffer (1.7 M sucrose, 20 mM Tris-HCl (pH 8.0), 2 mM MgCl_2_, 0.2% Triton X-100, 2 mM EDTA (pH 8.0), 15 mM β-Mercaptoethanol, 0.1 mM PMSF) in a 15 ml Falcon tube and then centrifuged at 2200 g for 20 min at 4 °C. The pellets were washed with 500 μl 1× NEB2 buffer (NEB, #B7202) for two times at 5000 rpm for 5 min at 4 °C and resuspended in 362 μl 1× Dpn II buffer (NEB, #R0543). For the permeabilization of nuclei, 38 μl 1% SDS was added to the nuclei, and the mixture was incubated at 65 °C for 10 min exactly. Then the tube was put on ice and 44 μl 10% Triton-X 100 was added to quench the SDS, avoiding the generation of bubbles. For chromatin digestion, 100 U (10 μl) Dpn II (NEB, #R0543) was added to the tube and incubated overnight at 37 °C on a rocking platform.

On the next day, the digestion reaction was stopped at 65 °C, 20 min. Then 15 μl T4 ligase (NEB, #M0202L) was added directly after the heat inactivation of Dpn II and incubate at 16 ℃ for 8 h for proximal ligation *in situ*. Next, Proteinase K (NEB, #P8107) was added at 1/10 volume and incubated at 65 ℃ overnight to reverse crosslink the nuclei. The DNA was purified with ZYMO DNA Clean & Concentrator kit (ZYMO, #D4014). 0.5× DNA Clean Beads (Vazyme, #N411-02) was used to select the DNA fragments larger than 1 kb. Then the DNA was subjected to Nanopore sequencing as previously described (Jia et al., 2020)

**Pore-C and Hi-C data analysis**

The FAST5 files containing Nanopore raw signals were basecalled using Guppy (v4.0.0+) with default parameters. Reads with quality-score less than 7 were filtered out. Quality check was done with Nanoplot (GNU General Public License v3.0) (De Coster et al., 2018). After preprocessing, both Pore-C libraries were feed to Pore-C tools ([Pore-C-Snakemake](https://github.com/nanoporetech/Pore-C-Snakemake), version 0.3.0) (Ulahannan et al., 2019). Pore-C pipeline including steps of reference genome virtual digestion, mapping (bwasw, default parameters, used TAIR10 as the reference genome), alignment filter, fragment deposit and thus get the multi-way contact related to each read (Supplementary Figure1). The multi-way contact (both direct and indirect) can be spear into pairwise contact as depicted in Supplemental Figure 1, which make Pore-C to generate more valid pairwise contact with similar counts of reads compared with Hi-C. The positions of KEE regions are based on a previous study (Grob et al., 2014).

The raw data of Hi-C (NCBI BioProject Accession: PRJNA227546, Run: SRR1029605, Wang et al., 2015) were processed following the 4DN projects Hi-C processing pipeline (Dekker et al., 2013). Pairwise interaction matrixes of Hi-C and Pore-C were normalized with cooler zoomify –balance (Abdennur and Mirny, 2020). The contact frequency was calculated by cooltools with logarithmic binning and normalized by the maximum value. All the 2D contact matrix heatmap in this article were plotted in log-scale (norm=LogNorm, vmax=50_000) with cooltools on balanced matrix (resolution=50 kb). Detail of the commands and parameters can be found in Supplementary Table 2.

**High-order interaction significance analysis**

To test the enrichment of multi-way interaction of KEEs, we compared the contact counts within KEEs and within 100 randomly selected control regions. According to their location on chromosomes (arm or pericentromere), we randomly chose the corresponding regions of each KEEs as the control. For two interacting sites that are located in the same chromosomes side (e.g. KEE3 and KEE4), control regions were selected based on both location and the fixed distance of the first site, like the “k-mer” concept adopted in SPRITE analysis (Quinodoz et al., 2018). The count of reads containing the interactions among KEEs was adopted as the observed value, and that for control sites served as the expected value. Mann-Whitney U test was adopted to calculate the *P-*values.

**Methylation analysis**

The CG methylation was called from Pore-C library with deepsignal-plant (v0.1.2) (Ni et al., 2021). Weighted methylation with bin size 100 bp was calculated for comparison with WGBS results (Schultz et al., 2012; Stroud et al., 2012), and weighted methylation with bin size 10 kb was calculated to show the whole genome methylation level. The reads of WGBS library GEO Accession GSE38286 (Stroud et al., 2012) were mapped to TAIR10 reference genome using BSMAP v2.90 (Xi and Li, 2009) with 8% mismatches (Hu and Wang, 2021; Schultz et al., 2012). The DNA methylation level of 100 bp bin was calculated based on the read count of methylated cytosines and unmethylated cytosines, as #C/(#C+#T) (Schultz et al., 2012; Stroud et al., 2012).

Pearson correlation coefficient (PCC) of DNA methylation level between Pore-C contacts and enrichment [analysis](javascript:;) was done as previously described (Li et al., 2019). 10000 Pore-C reads were randomly selected for this analysis. DNA methylation levels on Pore-C reads were identified using nanopolish (v 0.13.3) to keep the raw CG methylation information in chimeric reads (Simpson et al., 2017). Single fragment methylation levels were calculated as the fraction of CG methylation sites (Schultz et al., 2012), and only fragments that have at least two CpG sites were kept. To avoid the impact of interactions within short genomic distance, which might share similar chromosome states with alike epigenetic modifications that impair the independent analysis of contact and methylation, contacts with genomic distance over 25 kb, 100 kb and 1 Mb were selected from pair-wise interactions. PCC of DNA methylation were calculating between the Pore-C contacts. The methylation levels of pair contacts were shuffled independently and join pairs for calculation of the expected PCC. Significance analysis between the observed PCC and the expected were done with Fisher's r-to-Z transformation implemented in R package ‘cocor’ (Diedenhofen and Musch, 2015). All commands and parameters used in this study can be referred to Supplemental Table 2.

**Reference**

Abdennur, N. and Mirny, L.A. (2020) Cooler: scalable storage for Hi-C data and other genomically labeled arrays. *Bioinformatics* **36**, 311-316.

De Coster, W., D’Hert, S., Schultz, D.T., Cruts, M. and Van Broeckhoven, C. (2018) NanoPack: visualizing and processing long-read sequencing data. *Bioinformatics* **34**, 2666-2669.

Dekker, J., Marti-Renom, M.A. and Mirny, L.A. (2013) Exploring the three-dimensional organization of genomes: interpreting chromatin interaction data. *Nature Reviews Genetics* **14**, 390-403.

Diedenhofen, B. and Musch, J. (2015) cocor: A Comprehensive Solution for the Statistical Comparison of Correlations. *PloS one* **10**, e0121945.

Feng, S., Cokus, Shawn J., Schubert, V., Zhai, J., Pellegrini, M. and Jacobsen, Steven E. (2014) Genome-wide Hi-C Analyses in Wild-Type and Mutants Reveal High-Resolution Chromatin Interactions in Arabidopsis. *Molecular cell* **55**, 694-707.

Grob, S., Schmid, M.W. and Grossniklaus, U. (2014) Hi-C analysis in Arabidopsis identifies the KNOT, a structure with similarities to the flamenco locus of Drosophila. *Molecular cell* **55**, 678-693.

Hu, M. and Wang, S. (2021) Chromatin Tracing: Imaging 3D Genome and Nucleome. *Trends in cell biology* **31**, 5-8.

Jia, J., Long, Y., Zhang, H., Li, Z., Liu, Z., Zhao, Y., et al. (2020) Post-transcriptional splicing of nascent RNA contributes to widespread intron retention in plants. *Nature plants* **6**, 780-788.

Krzywinski, M., Schein, J., Birol, İ., Connors, J., Gascoyne, R., Horsman, D., et al. (2009) Circos: An information aesthetic for comparative genomics. *Genome research* **19**, 1639-1645.

Li, G., Liu, Y., Zhang, Y., Kubo, N., Yu, M., Fang, R., et al. (2019) Joint profiling of DNA methylation and chromatin architecture in single cells. *Nature methods* **16**, 991-993.

Ni, P., Huang, N., Nie, F., Zhang, J., Zhang, Z., Wu, B., et al. (2021) Genome-wide Detection of Cytosine Methylations in Plant from Nanopore sequencing data using Deep Learning. *bioRxiv*, 2021.2002.2007.430077.

Quinodoz, S.A., Ollikainen, N., Tabak, B., Palla, A., Schmidt, J.M., Detmar, E., et al. (2018) Higher-Order Inter-chromosomal Hubs Shape 3D Genome Organization in the Nucleus. *Cell* **174**, 744-757.e724.

Schultz, M.D., Schmitz, R.J. and Ecker, J.R. (2012) Leveling the playing field for analyses of single-base resolution DNA methylomes. *Trends in Genetics* **28**, 583-585.

Simpson, J.T., Workman, R.E., Zuzarte, P.C., David, M., Dursi, L.J. and Timp, W. (2017) Detecting DNA cytosine methylation using nanopore sequencing. *Nature methods* **14**, 407-410.

Stroud, H., Hale, C.J., Feng, S., Caro, E., Jacob, Y., Michaels, S.D., et al. (2012) DNA Methyltransferases Are Required to Induce Heterochromatic Re-Replication in Arabidopsis. *PLoS genetics* **8**, e1002808.

Ulahannan, N., Pendleton, M., Deshpande, A., Schwenk, S., Behr, J.M., Dai, X., et al. (2019) Nanopore sequencing of DNA concatemers reveals higher-order features of chromatin structure. *bioRxiv*, 833590.

Xi, Y. and Li, W. (2009) BSMAP: whole genome bisulfite sequence MAPping program. *BMC bioinformatics* **10**, 232.

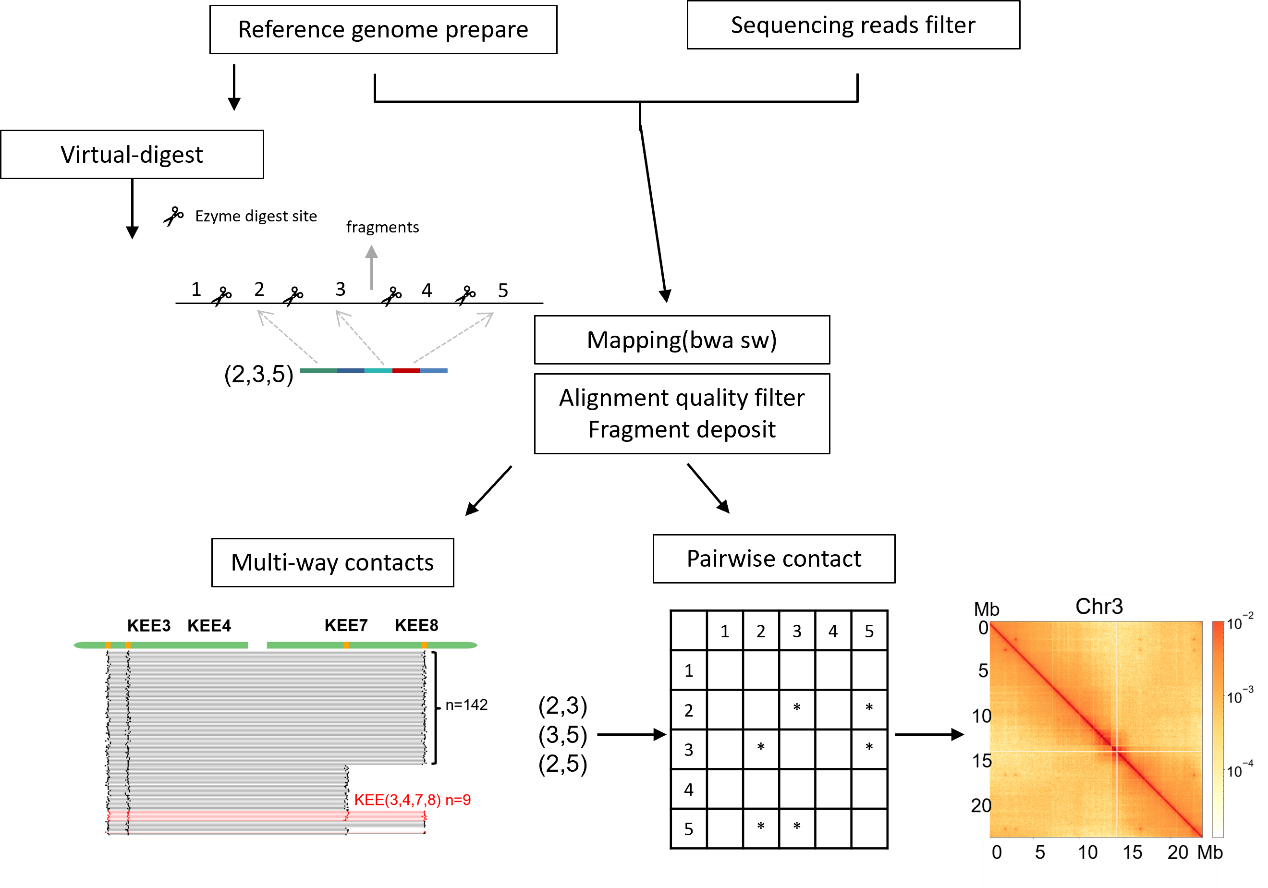

**Supplemental Figure 1 Description of Pore-C pipeline.**

After filtering by length and quality, Pore-C reads were aligned with reference genome. Virtual digestion was done on reference genome and alignment results were annotated accordingly. The interacting information can be used for multi-way interaction analysis or splited into pair-wise contact for contact matrix generation.

| **Supplemental Table 1. The sequence statistics of Pore-C libraries.** | | | |
| --- | --- | --- | --- |
| Sequence statistics | Pore-C Rep1 | Pore-C Rep2 | Pore-C merge |
| Mean read length (bp) | 1,811.8 | 2,262.10 | 1,970.20 |
| Mean read quality | 13.7 | 13.3 | 13.5 |
| Median read length (bp) | 1,326 | 1,743 | 1,481 |
| Median read quality | 13.9 | 13.5 | 13.8 |
| Number of reads | 6,452,571 | 3,499,745 | 9,952,316 |
| Read length N50 (bp) | 2,654 | 3,006 | 2,793 |
| STDEV read length (bp) | 1,557.0 | 1,781.1 | 1,653.4 |
| Total bases | 11,691,022,388 | 7,916,715,505 | 19,607,737,893 |
| Mapping rate | 96.50% | 97.04% | 96.70% |

**Supplemental Table 2. Commands and parameters used in this study**

|  | Steps | Software | Version | Mode | Parameters | Purpose | Reference |
| --- | --- | --- | --- | --- | --- | --- | --- |
| Nanopore sequence basecalling | step1 | Guppy | v4.0.0+ | / | --c dna_r9.4.1_450bps_hac.cfg | basecalling | / |
| Quality control | step1 | Nanoplot | v3.0 | / | default parameters | quality control | Coster, D' Hert, et al., 2018 |
| Pore-C data analysis | step1 | [Pore-C-Snakemake](https://github.com/nanoporetech/Pore-C-Snakemake) | 0.3.0 | / | default parameters | decode the interaction information in Pore-C reads | Ulahannan et al., 2019 |
| Hi-C data analysis | step1 | bwa | 0.7.17 | mem | -SP5M | mapping | Li H. 2013 |
|  | step2 | pairsamtools | 0.0.1-dev | parse | default parameters | generating pair contact information | / |
|  | step3 | pairtools | 0.3.0 | sort | default parameters |  | / |
|  | step4 | pairtools | 0.3.0 | dedup | default parameters | remove PCR duplicates | / |
|  | step5 | bgzip | 1.9 |  | default parameters |  |  |
|  | step6 | pairix | 0.3.7 |  | -f -p pairs |  | / |
|  | step7 | cooler | 0.8.7+ | cload pairix | default parameters | generating contact matrix | Abdennur and Mirny, 2020 |
|  | step8 | cooler | 0.8.7+ | zoomify | --balance --balance-args --convergence-policy=store_nan --balance-args --max-iters=10000 | generating balanced contact matrix with multiple resolution | Abdennur and Mirny, 2020 |
| Methylation genomic distribution analysis | / | deepsignal-plant | v0.1.2 | / | default | call CG, CHG, CHH methylation from Pore-C reads | Ni et al., 2021 |
| Methylation-Contact correlation analysis |  | nanopolish | v0.13.3 | / | -q cpg --min-mapping-quality=1 | call CG methylation from Pore-C reads(keep methyaltion information on chimeric reads) | Simpson et al., 2017 |
| WGBS methylation analysis |  | BSMAP | v2.90 | / | 8% mismatches | map methylation to reference genome | Yuanxin Xi and Wei Li, 2009 |
